# Identification of 2CS-CHX^T^ operon signature of chlorhexidine tolerance among *Enterococcus faecium*

**DOI:** 10.1101/704346

**Authors:** Bárbara Duarte, Ana P. Pereira, Ana R. Freitas, Teresa M. Coque, Anette M. Hammerum, Henrik Hasman, Patrícia Antunes, Luísa Peixe, Carla Novais

**Affiliations:** UCIBIO/REQUIMTE. Department of Biological Sciences. Laboratory of Microbiology. Faculty of Pharmacy. University of Porto. Porto. Portugal; Faculty of Nutrition and Food Sciences. University of Porto. Porto. Portugal; Servicio de Microbiologia. Hospital Universitario Ramón y Cajal. Instituto Ramón y Cajal de Investigación Sanitaria (IRYCIS), Madrid, Spain; Centro de Investigación Biomédica en Red de Epidemiología y Salud Pública (CIBER-ESP), Madrid, Spain; Unidad de Resistencia a Antibióticos y Virulencia Bacteriana asociada al Consejo Superior de Investigaciones Científicas (CSIC), Madrid, Spain; Department of Bacteria, Parasites and Fungi, Statens Serum Institut, Copenhagen, Denmark

**Keywords:** Chlorhexidine-tolerance, *Enterococcus faecium*, 2CS-CHX^T^ operon, epidemiological-cutoff

## Abstract

Chlorhexidine (CHX) is a broad-spectrum antiseptic widely used in community and clinical contexts for many years, recently acquiring higher relevance in nosocomial infections control worldwide. Despite of this, CHX tolerance has been poorly understood among *Enterococcus faecium*, one of the leading agents causing nosocomial infections. This study provides new phenotypic and molecular data for a better identification of CHX tolerant *E. faecium* subpopulations in community and clinical contexts. MIC_CHX_ distribution of 106 *E. faecium* suggested the occurrence of tolerant subpopulations in diverse sources (human, animal, food, environment) and phylogenomic backgrounds (clades A1/A2/B), with predominance in clade A1. They carried a specific variant of 2CS-CHX^T^ operon, here identified. It encodes a glucose and an amino-acid-polyamine-organocation family transporters, besides the DNA-binding-response-regulator ChtR with a P102H mutation previously described in only CHX tolerant clade A1 *E. faecium*, and the ChtS sensor. Combined data from normal MIC distribution and 2CS-CHX^T^ operon characterization supports a tentative epidemiological cut-off (ECOFF) of 8 mg/L to CHX, useful to detect tolerant *E. faecium* populations in future surveillance studies. The spread of tolerant *E. faecium* in diverse epidemiological backgrounds alerts for a prudent use of CHX in multiple contexts.

**Importance:** Chlorhexidine is one of the substances included in the World Health Organization’s List of Essential Medicines which comprises the safest and most effective medicines needed in global health systems. Although it has been widely applied as a disinfectant and antiseptic since the 1950s in healthcare (skin, hands, mouthwashes, eye drops), its use in hospitals to prevent nosocomial infections has increased worldwide in recent years. Here we provide a comprehensive study on chlorhexidine tolerance among *Enterococcus faecium*, one of the leading nosocomial agents worldwide, and identified a novel 2CS-CHX^T^ operon as a signature of tolerant strains occurring in diverse phylogenomic groups. Finally, our data allowed to propose a tentative epidemiological cut-off of 8 mg/L, useful to detect tolerant *E. faecium* populations in future surveillance studies in community and clinical contexts.

## INTRODUCTION

Chlorhexidine is one of the substances included in the World Health Organization’s List of Essential Medicines which comprises the safest and most effective medicines needed in global health systems (1). It is a cationic biguanide acting in bacteria by binding to cell wall and membrane anionic sites and affecting the osmotic equilibrium of the cell. At elevated concentrations, as used in antimicrobial marketed products, it causes the loss of membrane structural integrity, resulting in cell death (2, 3). Chlorhexidine is widely used as a topical antiseptic in the community (e.g., oral care, skin antisepsis, cosmetics), in veterinary (e.g., skin antisepsis, teat dips, udder and eye wash) and hospitals (e.g., hand hygiene, daily patient baths, impregnated wash cloths, impregnated cardiovascular catheters, oral antisepsis) with concentrations varying between 0.05-4% (corresponding to 500-40,000 mg/L) (3–6). Although chlorhexidine is in the market since the 1950’s, it has been increasingly used in the last decade in the hospital setting to prevent catheter-related bloodstream infections, ventilator associated pneumonia, skin soft infections or patients’ skin colonization by multidrug resistant bacteria included in high priority WHO list, as methicillin-resistant *Staphylococcus aureus* (MRSA) or vancomycin resistant *Enterococcus* (VRE) (4, 7–10, https://www.who.int/medicines/publications/WHO-PPL-Short_Summary_25Feb-ET_NM_WHO.pdf).

*Enterococcus faecium* is one of the species most commonly associated with nosocomial infections worldwide in the last years, with clinical isolates often being resistant to first-line antibiotics, ampicillin and vancomycin (11). Despite the high transmissibility of *E. faecium* strains in health care institutions, few studies have evaluated the chlorhexidine activity against this species. Most of them focused on *E. faecium* from a single source (either clinical, animal or food) (12–19), did not consider strain’s genetic background (12–14, 16, 17, 19, 20) or included a low number of isolates (14–17), precluding the analysis of chlorhexidine susceptibility data in the context of *E. faecium* population structure.

The genotypes or expression profiles of *E. faecium* isolates showing chlorhexidine tolerance have been scarcely explored (13, 21–23). Tolerance phenotypes were related to a unique, nonsynonymous single amino acid polymorphism (P102H) in a conserved DNA-binding response regulator ChtR in three strains from clade A1 of clinical origin, which along with a histidine kinase sensor ChtS formed a regulator system called 2CS with an unknown regulon. The advantage of ChtR-P102H mutation for *E. faecium* was demonstrated by molecular (mutants with ChtR/ChtS deletion and trans complementation) and phenotypic studies involving growth curves in the presence of 1.2 mg/L of chlorhexidine (21). Also, the occurrence of A290V amino acid change in EfrE protein (codifying for predicted ABC transport system) detected in mutants generated after 21 days of passages in culture media supplemented with sub-inhibitory chlorhexidine concentrations was again detected in a clade A1 *E. faecium* (wild type: MIC=4,9 mg/L; mutants: MIC=19,6 mg/L) (23). Among Gram-positive bacteria, cross-tolerance to quaternary ammonium compounds and chlorhexidine was suggested to be associated to *qacA/B* gene in *Staphylococccus* spp. (24, 25), with the limited data for *E. faecium* (17, 26) impairing to assess the potential role of this or other *qac/smr* genes in chlorhexidine tolerance among this species.

This study provides new phenotypic and molecular data for a better identification of chlorhexidine tolerant *E. faecium* subpopulations in community and clinical contexts. It is demonstrated that tolerant *E. faecium* are spread in different sources and phylogenomic groups with predominance in clade A1, and that they carry a genetically stable 2CS-CHX^T^ operon, here identified. The combined phenotypic and molecular data points for a chlorhexidine tentative ECOFF of 8 mg/L for *E. faecium*, that could be used in future surveillance studies to detect tolerant populations, both in community and clinical contexts.

## RESULTS

### Chlorhexidine susceptibility of *E. faecium* from different phylogenomic groups and sources

Chlorhexidine MIC (MIC_CHX_) among the 106 *E. faecium* tested ranged between ≤2 to 32 mg/L, with a mode in the 16 mg/L (Fig. 1-I; Table 1). The ECOFF for 95% of the population suggested by the ECOFFinder tool was ≤32mg/L. However, the MIC_CHX_ distribution presented a selected log2 SD=1.17 and a fitted curve slightly deviated to the left comparing to raw data distribution, suggesting a tolerant subpopulation in the *E. faecium* sample (Fig. 1). Also, the NORM.DIST Excel function indicates that the probability of an isolate having a MIC_CHX_>16mg/L is of 0%, suggesting that at least those with MIC_CHX_=32mg/L have an acquired tolerance phenotype. Although ECOFF are not determined for species subpopulations, in order to evaluate a potential variability among different *E. faecium* clonal lineages and the potential occurrence of tolerant isolates within each clade, *E. faecium* from clades A1, A2 and B were independently analysed. A unimodal distribution was observed for *E. faecium* from both clades A1 (Fig.1-II) and B (Fig.1-IV), with those from clade A1 presenting a mode at 16 mg/L (77% of the 48 isolates of clade A1) and from clade B a mode at 8 mg/L (60% of the 15 isolates of clade B) (Table 1; Fig. 1). The upper limit for 95% of the population suggested by the ECOFFinder tool was of 16 mg/L for both clades. However, the probability of occurrence of a wild-type isolate with MICCHX>8 and ≤16 mg/L in clade A1 calculated by the NORM.DIST Excel function was of 99.9% and in clade B of 0.00014%. In both cases, the probability of occurrence of an isolate with MIC_CHX_>16 mg/L was of 0%. Contrasting with these data, the distribution of *E. faecium* from clade A2 was bimodal (modes at 4 mg/L-30% and 16 mg/L-37% of 43 isolates of clade A2), suggesting the occurrence of two subpopulations (Fig. 1-IIIa). The limit of 95% of wild type population of the clade A2 was proposed as 8 mg/L by the ECOFFinder tool with the probability of a wildtype isolate to have a MIC_CHX_>4 and ≤8mg/L being of 2.1% and MIC>8 mg/l of 0%. The analysis of the distribution data and the probability of occurrence of wild type isolates in diverse MIC_CHX_ of clades A2 and B suggests a tentative ECOFF of 8 mg/L for *E. faecium*.

**FIG 1.**
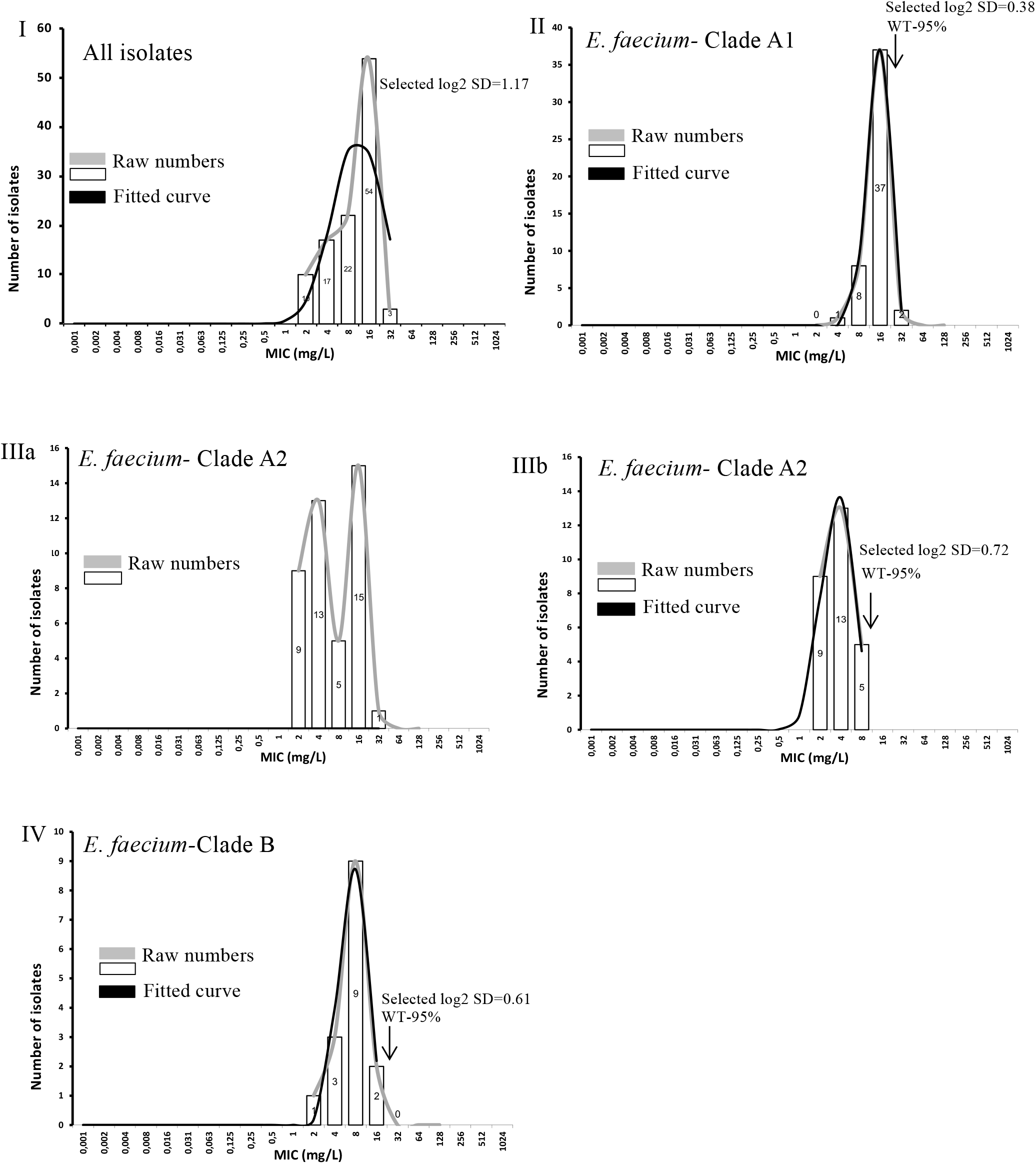
Distribution of the *Enterococcus faecium* studied by different chlorhexidine MICs. The graphs fitted curves and limits of 95% of wild type population (L95-WT) were drawn based on the calculations of ECOFFINDER tool. Graph I: distribution of the 106 *E. faecium* studied (all clades). Graph II: distribution of clade A1 *E. faecium* (n=48; L-WT= ≤16 mg/L). Graph IIIa: distribution of raw data of clade A2 *E. faecium* (n=43). Graph IIIb: distribution of clade A2 *E. faecium* (n=27) without isolates presenting MICs values corresponding to the second mode and higher. The removal of such isolates is necessary for ECOFFINDER tool be able to propose the L95-WT, in this case ≤8 mg/L. Graph IV: distribution of clade B *E. faecium* (n=15; L-WT≤16 mg/L). The DIST.NORM Excel’s-16.19 function indicates that in all cases the probability of occurrence of an isolate with a MIC>16 mg/L is 0% and variable among clades in the range MIC>8 and ≤ 16mg/L: 99,9% for clade A1, 0% for clade A2 and 0.00014% for clade B. MIC=2mg/L represents MIC≤2 mg/L.

**TABLE 1.**
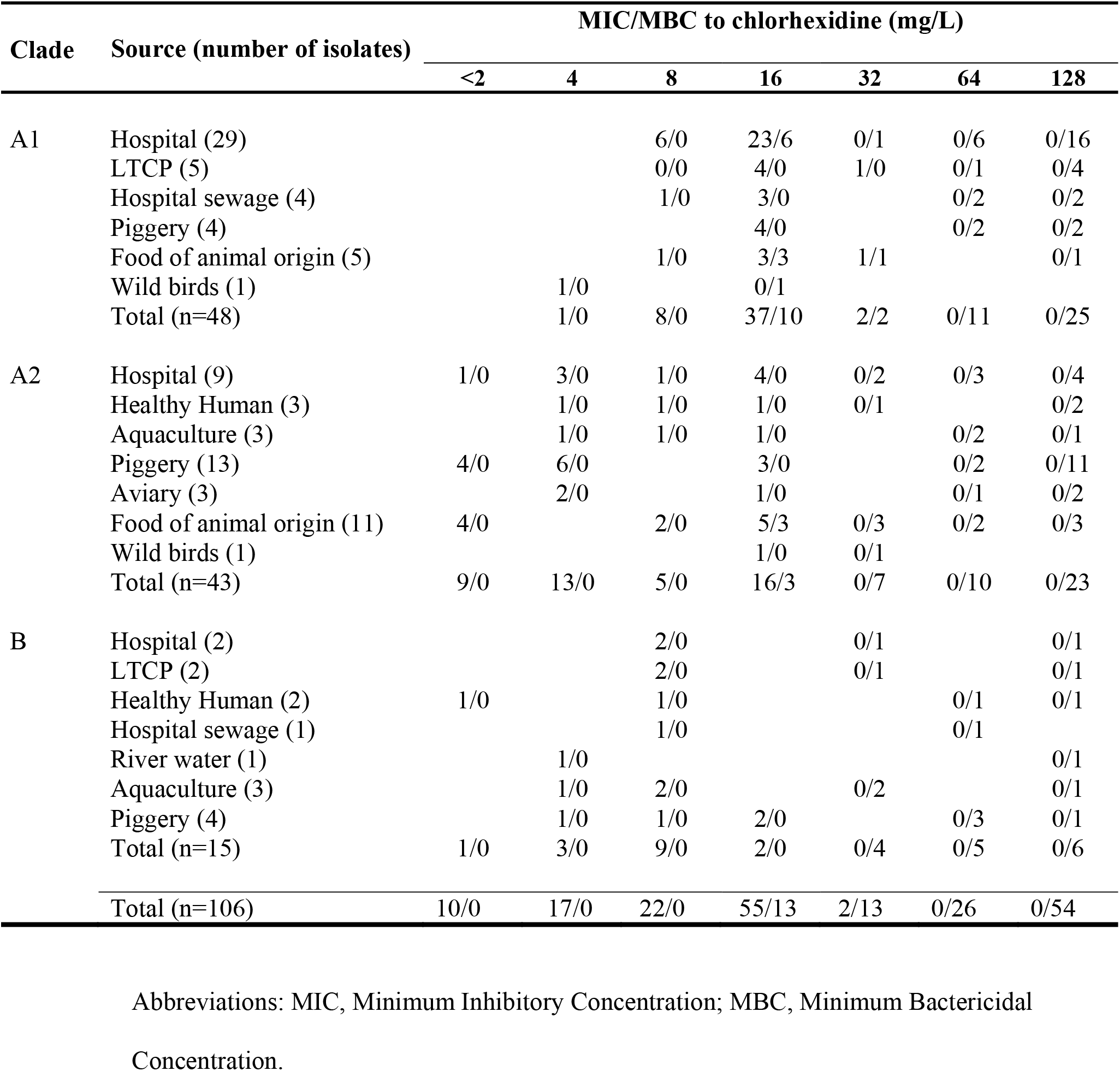
Minimum inhibitory concentration and minimum bactericidal concentration of chlorhexidine to *Enterococcus faecium* from different clades and sources.

| Clade | Source (number of isolates) | MIC/MBC to chlorhexidine (mg/L) |  |  |  |  |  |  |
| --- | --- | --- | --- | --- | --- | --- | --- | --- |
|  |  | <2 | 4 | 8 | 16 | 32 | 64 | 128 |
| A1 | Hospital (29) |  |  | 6/0 | 23/6 | 0/1 | 0/6 | 0/16 |
|  | LTCP (5) |  |  | 0/0 | 4/0 | 1/0 | 0/1 | 0/4 |
|  | Hospital sewage (4) |  |  | 1/0 | 3/0 |  | 0/2 | 0/2 |
|  | Piggery (4) |  |  |  | 4/0 |  | 0/2 | 0/2 |
|  | Food of animal origin (5) |  |  | 1/0 | 3/3 | 1/1 |  | 0/1 |
|  | Wild birds (1) |  | 1/0 |  | 0/1 |  |  |  |
|  | Total (n=48) |  | 1/0 | 8/0 | 37/10 | 2/2 | 0/11 | 0/25 |
| A2 | Hospital (9) | 1/0 | 3/0 | 1/0 | 4/0 | 0/2 | 0/3 | 0/4 |
|  | Healthy Human (3) |  | 1/0 | 1/0 | 1/0 | 0/1 |  | 0/2 |
|  | Aquaculture (3) |  | 1/0 | 1/0 | 1/0 |  | 0/2 | 0/1 |
|  | Piggery (13) | 4/0 | 6/0 |  | 3/0 |  | 0/2 | 0/11 |
|  | Aviary (3) |  | 2/0 |  | 1/0 |  | 0/1 | 0/2 |
|  | Food of animal origin (11) | 4/0 |  | 2/0 | 5/3 | 0/3 | 0/2 | 0/3 |
|  | Wild birds (1) |  |  |  | 1/0 | 0/1 |  |  |
|  | Total (n=43) | 9/0 | 13/0 | 5/0 | 16/3 | 0/7 | 0/10 | 0/23 |
| B | Hospital (2) |  |  | 2/0 |  | 0/1 |  | 0/1 |
|  | LTCP (2) |  |  | 2/0 |  | 0/1 |  | 0/1 |
|  | Healthy Human (2) | 1/0 |  | 1/0 |  |  | 0/1 | 0/1 |
|  | Hospital sewage (1) |  |  | 1/0 |  |  | 0/1 |  |
|  | River water (1) |  | 1/0 |  |  |  |  | 0/1 |
|  | Aquaculture (3) |  | 1/0 | 2/0 |  | 0/2 |  | 0/1 |
|  | Piggery (4) |  | 1/0 | 1/0 | 2/0 |  | 0/3 | 0/1 |
|  | Total (n=15) | 1/0 | 3/0 | 9/0 | 2/0 | 0/4 | 0/5 | 0/6 |
| Total (n=106) |  | 10/0 | 17/0 | 22/0 | 55/13 | 2/13 | 0/26 | 0/54 |

Abbreviations: MIC, Minimum Inhibitory Concentration; MBC, Minimum Bactericidal Concentration.

Minimum bactericidal concentrations of chlorhexidine (MBC_CHX_) ranged between 16-128mg/L with MBC95 and MBC99=128 mg/L (Table 1). The highest MBC_CHX_ of 128 mg/L was observed in 51% (n=54/106) of the isolates studied, belonging to all clades, diverse sources and presenting MIC_CHX_<2-32mg/L. Of note, isolates associated with human infections (n=40; clades A1, A2, B) showed the same MBC95 and MIC99=128 mg/L than those not associated with human infections (n=66; clades A1, A2, B). When microdilution assays were performed at pH=5, MBC_CHX_ values increased 1- to 4-fold dilutions for 32 isolates out of 37 tested comparing to MBC_CHX_ in standard conditions. For 27 *E. faecium* the MBC_CHX_ increased to 256 mg/L (Table 2).

**TABLE 2.**
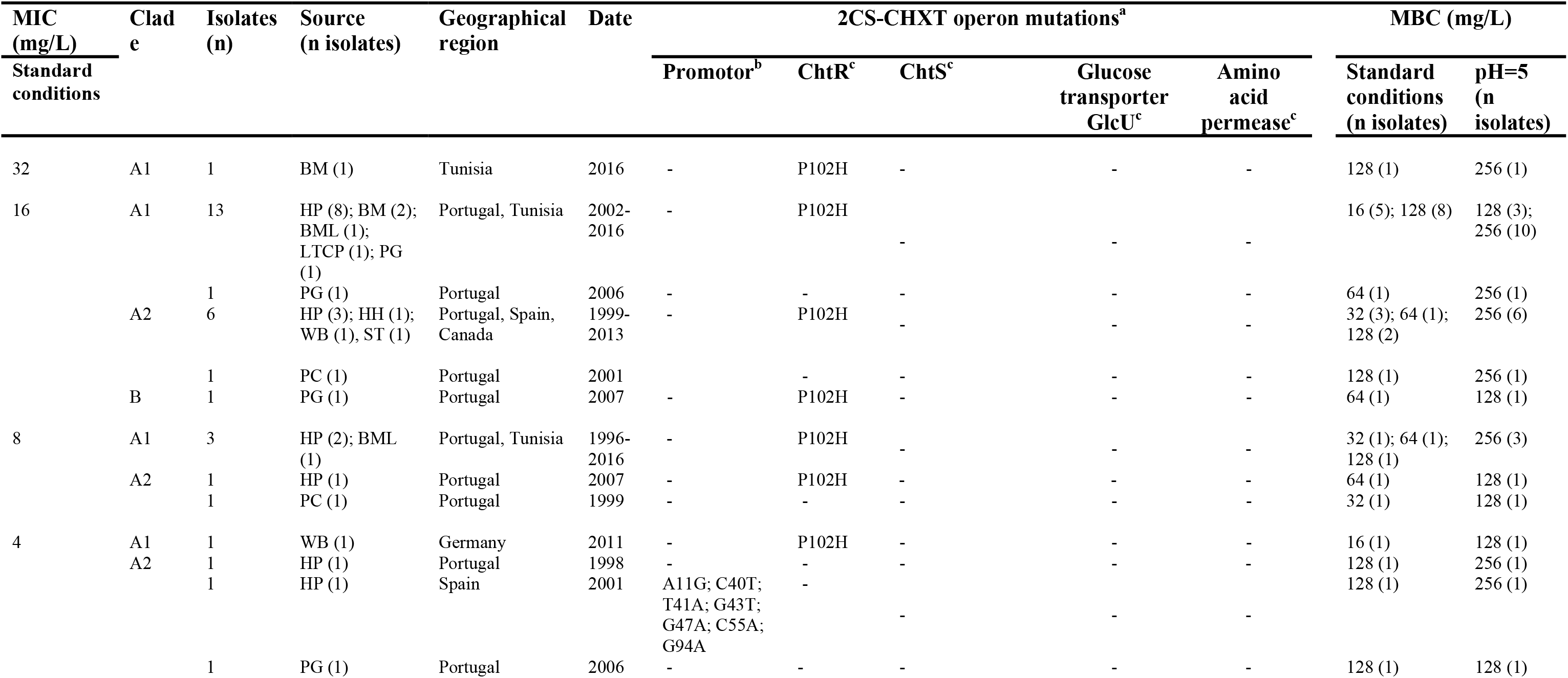
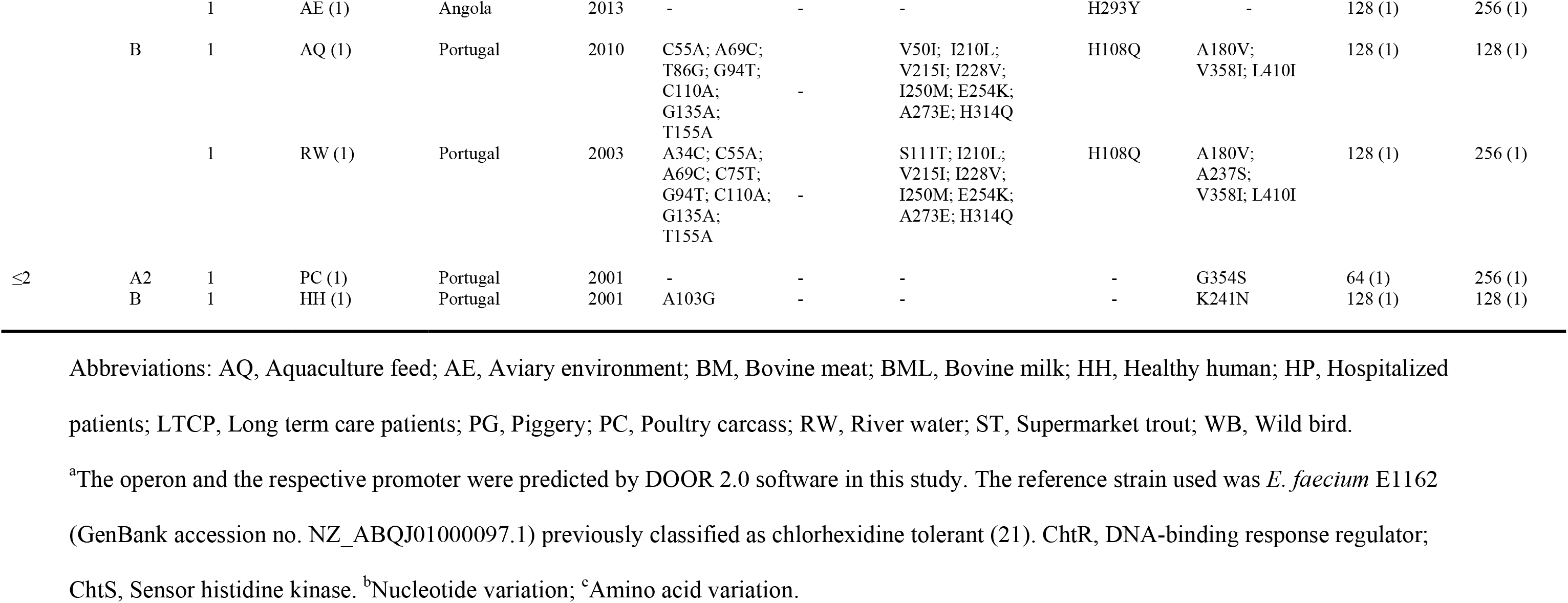
Distribution of the 2CS-CHX^T^ operon variants among *E. faecium* isolates showing different chlorhexidine phenotypes and with diverse epidemiological backgrounds.

### Chlorhexidine susceptibility of *E. faecium* with different antibiotic resistance profiles

Ampicillin resistant *E. faecium* presented higher MIC99_CHX_ than ampicillin susceptible isolates (32mg/L vs 16 mg/L), although such a difference was determined by only two isolates. The MIC95_CHX_, MBC95_CHX_ and MBC99_CHX_ were the same for ampicillin resistant and susceptible *E. faecium* (16mg/L, 128 mg/L and 128mg/L, respectively) (Table S2). No differences were detected among MIC95_CHX_ and MIC99_CHX_ (16mg/L and 32 mg/L, respectively) and MBC95_CHX_ and MBC99_CHX_ (128 mg/L in both cases) when comparing vancomycin-resistant and vancomycin-susceptible *E. faecium*.

### Distribution of the ChtR-P102H mutation in *E. faecium* from different phylogenomic groups and sources

The P102H mutation of the ChtR protein sequence, previously described to be associated with clorhexidine tolerance (21), was detected in 26 of the 37 (70%) genomes studied here, which were recovered from isolates of disparate sources (hospitalized patients, n=14; patient at long-term-care-facility, n=1; healthy human, n=1; piggeries, n=2; food of animal origin, n=6; wild birds, n=2) and phylogenomic groups (clades A1: 69%, n=18; A2: 27%, n=7; B: 4%, n=1). Isolates without such mutation were mostly from non-clinical origins (n=9 out of 11) and belong to clades A2 (64%, n=7; including two clinical isolates), B (27%, n=3) and A1 (9%, n=1). Similar results were found among the 794 genomes analysed and published at the GenBank database (Tables S3 and S4). It was verified that 71% (n=562/794) presented the ChtR-P102H mutation, mostly associated with the clade A1 (79%, n=445/562), followed by clade A2 (20%, n=113/562) and clade B (1%, n=4/562) (p<0.05; Chi square test; GraphPad Prism version 8.0). Among the 232 genomes without the ChtR-P102H, most were from clade A2 (73%, n=170/232) followed by clade B (18%, n=42/232) and clade A1 (9%, n=20/232).

### Identification of the operon regulated by the ChtR protein and its association with chlorhexidine tolerance phenotype

The DOOR software predicted that the *chtR* gene was part of an operon with a length of 4086 bp, containing a histidine kinase sensor (*chtS*), previously described by Guzmán-Prieto et al (21), and genes coding for proteins related to a glucose:proton symporter GlcU [superfamily (cl26915) of second carriers of sugar transport transmembrane proteins] and an amino acid permease of the Amino acid-Polyamine-organoCation (APC) family [superfamily (cl26159) AA_permease_2; GabP type transporters working as GABA and other amino acid a H+ symporters], here identified. The operon was located between 59 678 nt and 63 763 nt positions of the *E. faecium* E1162 reference strain (GenBank accession no. NZ_ABQJ01000097.1). The predicted operon promoter corresponded to the sequence between 63946-63779 nt position from E1162 reference strain (GenBank accession no. NZ_ABQJ01000097.1), distancing 15 nucleotides from the *chtR* gene. From now on this operon will be called as 2CS-CHX^T^ by adapting the previous designation used by Guzmán-Prieto et al (21) for the regulator and sensor genes.

The MIC_CHX_ of isolates carrying the ChtR-P102H mutation ranged from 4 to 32 mg/L, with 81% of them showing a MIC_CHX_≥16 mg/L (14 from clade A1, 6 from clade A2, 1 from clade B), followed by 15% with MIC_CHX_=8 mg/L (3 from clade A1, 1 from A2) and 4% with MIC_CHX_=4mg/L (1 isolate from clade A1). The isolates without the P102H mutation presented MIC_CHX_ ranging from <2 to 16 mg/L, with 73% (n=8/11; 5 from clade A2 and 3 from B) showing a MIC_CHX_≤4 mg/L, 18% (n=2/11; 1 from each clade A1 and A2) a MIC_CHX_=16 mg/L and 9% (n=1/11; clade A2) a MIC_CHX_=8 mg/L (Table 2). These results point for a higher ChtR-P102H occurrence in isolates with a MIC_CHX_≥8mg/L (p<0.05; Fisher exact test; GraphPad Prism version 8.0). The analysis of the sequences of the 37 *E. faecium* genomes from this study showed that those carrying the P102H mutation in the ChtR (n=26) presented a 2CS-CHX^T^ operon sequence identity with the reference strain *E. faecium* E1162 (Table 2), and all but one isolate showed a MIC_CHX_≥8 mg/L. A higher variability was observed among those 2CS-CHX^T^ operons not carrying the ChtR-P102H mutation, mostly on those isolates with a MICCHX≤4mg/L. In this group, diverse mutations in promoter, sensor histidine kinase, glucose transporter or amino acid permease were observed (Table 2). No clear differences were observed in MBC_CHX_ among isolates carrying or not the ChtR-P102H mutation, both in standard and pH=5 adjusted conditions.

### Detection of A290V mutation in EfrE protein and the occurrence of *qac/smr* genes

The A290V mutation in EfrE protein was not detected in any isolated analysed, even on those isolates with MIC_CHX_>16 mg/L, contrasting with laboratory generated isolates with this phenotype (23). This mutation was also absent in the selected 980 *E. faecium* genomes published at the GenBank database with EfrR protein identified.

A hospital sewage *E. faecium* carrying *qacZ* gene (27) presented a MIC/MBC=8/64 mg/L to chlorhexidine similarly to other isolates without such gene. The analysis of the 37 *E. faecium* genomes did not reveal the occurrence of any of the *qac/smr* genes variants searched in this study.

## Discussion

The increasing use of chlorhexidine in recent years foresees an evolution of bacterial populations towards tolerance, imposing the need to identify chlorhexidine wild type *E. faecium* populations to better establish epidemiological cutoffs (ECOFFs) for susceptibility surveillance studies. An ECOFF (MIC) to chlorhexidine of ≤32 mg/L was recently proposed by Morrissey et al. (20). However, our analysis of these authors’ results by using the ECOFFinder tool (data not shown) suggests the presence of a tolerant population by a fitted curve slightly deviated to the left comparing to raw data distribution, as in the case of our 106 isolates, and thus an overestimation of the proposed ECOFF. The bimodal distribution of *E. faecium* within clade A2 in our study also supports the presence of tolerant populations. The occurrence probability of 0% or almost 0% (NORM.DIST function) of *E. faecium* with MIC_CHX_>8 mg/L within clades A2 or B, respectively, suggests a tentative ECOFF of 8mg/L, although it classifies most of clade A1 isolates as tolerant (81%; MIC=16-32mg/L). Clade A1 subpopulation is predominant in the hospital setting where the great use of chlorhexidine could be favouring their evolution towards tolerance to this antiseptic, as occurred among *Staphylococcus* spp. causing human infections (28, 29). A bias imposed by the high number of isolates from clade A1 in our study, could have influenced the initial proposal of the ECOFFinder tool of an ECOFF=32 mg/L for the 106 *E. faecium* as well as the ECOFF=32 mg/L proposed in Morrissey et al. (20) study, in which the *E. faecium* genetic background was not described. Such bias might also justify variable data among studies associating chlorhexidine tolerance with vancomycin resistance and not considering strains genetic background (16, 18, 19). In fact, ampicillin resistance, and not vancomycin, is an important marker of clade A1 strains (30), which explains the higher MIC99_CHX_ among *E. faecium* resistant to ampicillin observed in our study. The analysis of genotypes associated with chlorhexidine tolerance showed that ChtR-P102H mutation is mostly present in isolates with a MIC_CHX_≥8 mg/L from any clade and recovered from different sources, contrasting with previous descriptions only in clade A1 *E. faecium* (21). The more antique strains with ChtR-P102H mutation found in available genomes were from clade A2 and recovered from milk and dairy utensils (USA, 1927), a mouse (The Netherlands, 1959), human clinical isolates (The Netherlands, 1960-1965) and others of non-identified source and origin (≤1946). Data from our and published genomes suggest that independent ChtR-P102H mutation acquisition could have occurred in isolates from different genetic and epidemiological backgrounds. However, the main presence of ChtR-P102H in isolates from clade A1, again supports the adaptation of this subpopulation towards chlorhexidine tolerance. Clade A1 strains from 1987 (UK) and early 1990’s (USA, France) are the oldest and within the time frame of the emergence of this population in the hospital setting (30). Nevertheless, a careful analysis is necessary, as bias associated with published genomes should be considered.

The new genes here predicted to be regulated by the ChtR and to complete the new 2CS-CHX^T^ operon code for a second carrier symport glucose/H+ transporter (GlcU type), and a permease of the APC family, which seem to be associated with gamma-aminobutyric acid (GABA) or other amino acids transport. The GlcU is widespread in Firmicutes (31) and is a member of the prokaryote glucose/ribose porter family, belonging to the drug metabolite superfamily of transporters (32). The one found in *E. faecium* has 52% of amino acidic identity with those of *Lactococcus lactis* M1363 (GenBank: CAL99123.1), described to be a glucose uptake transporter dependent of proton-motive force, which is preferentially expressed when environmental conditions inhibit the high affinity-glucose uptake systems sugar phosphoenolpyruvate-phosphotransferases systems (PEP-PTS) (31). In different Firmicutes low concentrations of chlorhexidine inhibit the PEP-PTS system (31, 33–36). Among *Streptococcus* spp. sugar metabolism continues by a sugar-uptake system driven by proton motive force insensitive to this antiseptic (33, 34), which seems to have a higher expression at high rates of bacterial growth or acidic pH (37). Assuming that chlorhexidine inhibits PEP-PTS systems in *E. faecium* as in other Firmicutes and considering that the chlorhexidine susceptibility assays were performed in the exponential phase when a high-rate growth occurs, it seemed that the GlcU transporter could have an important role in the uptake of glucose for the bacterial cell survival under chlorhexidine stress. Finally, the amino acid permease was identified as belonging to the APC family, type GabP, which in some bacteria is induced by nitrogen limited culture conditions (38). We hypothesize that 2CS-CHX^T^ operon could contribute to a more effective energy associated metabolism used by the cell to cope with sub-inhibitory concentrations of chlorhexidine, although functional studies associated with each gene of the operon are still needed. It is of note that every time ChtR-P102H mutation is present, the 2CS-CHX^T^ sequence was the same among isolates from all clades, contrasting with the variable mutations in all operon genes and/or the promotor in *E. faecium* without the ChtR-P102H and with a MIC_CHX_≤4mg/L. The potential fixation of a stable 2CS-CHX^T^ operon with the ChtR-P102H mutation in isolates with higher MICCHX values suggests a role of this operon for chlorhexidine tolerance and that it could have contributed to the adaptation of particular subpopulations, as clade A1 in hospitals. The 2CS-CHX^T^ with the ChtR-P102H mutation could also have importance in other stress contexts, still unexplored, as pointed by its wide distribution as well as occurrence in strains collected before the introduction of chlorhexidine, which was commercialized in 1954.

In our study the acid pH=5 increased the MBC_CHX_ in several isolates. However, the lack of correlation between this higher MBC and 2CS-CHX^T^ variants suggests that other mechanisms of tolerance to stress, namely in response to acid pH, might contribute to higher survival of *E. faecium* to chlorhexidine in different conditions and contexts (e.g. survival of particular strains in the skin with pH=4-6) (39). Cross tolerance between acidic pH (pH=5.5) and chlorhexidine was also described to some *Streptococcus mutans* mutants, which seemed to be associated to changes in membrane fatty acids profiles (40).

The absence of the A290V mutation in the EfrE protein in all genomes analysed (our 37 *E. faecium* and available at GenBank database) indicates that A290V is not expected to often occur in wild type strains, namely in those with higher MIC/MBC_CHX_ (23). Moreover, the MIC/MBC_CHX_ values observed were not correlated with the presence of *qac/smr* genes analysed among *E. faecium*.

In conclusion, the previously proposed ECOFF (MIC) of 32mg/L for chlorhexidine (20) seems to be overestimated for *E. faecium* populations and our analysis of MIC_CHX_ normal distribution suggest a tentative ECOFF of 8 mg/L. However, the occurrence of the ChtR-P102H in *E. faecium* with MIC_CHX_=8mg/L, of few isolates without such mutation but with high MIC_CHX_ values and the lack of correlation between MBCs and 2CS-CHX^T^ variants underline the need of more studies to better understand *E. faecium* chlorhexidine tolerance mechanisms. The data also suggest that *E. faecium* from clade A1 is a more evolved population towards chlorhexidine tolerance, probably adapted to the high use of this antiseptic in the clinical setting. Independently of *E. faecium* clades or antibiotic susceptibility profile, our and other author’s data show that *E. faecium* have lower MBC_CHX_ (e.g. 256 mg/L) than concentrations used in marketed biocidal products (e.g. 4% chlorhexidine corresponds to 40,000mg/L) (12, 14, 18, 41). However, the data could still be of concern due to the low chlorhexidine concentrations occurring in different environments (e.g. residual skin concentrations are variable according to chlorhexidine baths procedures: 18.75-387.1 mg/L; sewage: <1 mg/L) (42–44), or bacteria could better adapt in specific conditions (e.g. acid pH, starvation) (40, 45), increasing their probability of survival under chlorhexidine stress. As in the case of other antimicrobials (e.g. antibiotics, ethanol) (46), a prudent use of chlorhexidine (e.g. standard protocols with sufficient exposure times and concentrations; control of its release to the environment) is important to maintain its efficacy.

## MATERIALS AND METHODS

### Bacterial isolates epidemiological background

One hundred and six *E. faecium* isolates obtained in Portugal, Spain, Angola, Canada, Germany, and Tunisia (1995-2016) during previous surveillance studies were included (11, 47–52) (Table S1). The clonal relationship of the 106 isolates was established by MLST, goeBURST and Bayesian Analysis Population Structure (BAPS) schemes (30, 53, http://pubmlst.org). They corresponded to 52 known sequence types (ST) and 13 BAPs subgroups clustering into clades A1 [n=48; hospitalized patients, n=29 isolates; long-term-care patients, n=5; hospital sewage, n=4; animal production setting (piggery), n=4; food of animal origin (bovine meat and milk), n=5; wild birds (rook), n=1], A2 [n=43; hospitalized patients, n=9; healthy humans, n=3; animal production setting (piggery, aquaculture, aviary), n=19; food of animal origin (poultry carcass, trout), n=11; wild birds (crow), n=1] and B [n=15; hospitalized patients, n=2; long-term-care patients, n=2; healthy humans, n=2; aquatic environment (hospital sewage, river water), n=2; animal production setting (piggery, aquaculture), n=7]. These isolates comprise 72% (n=76/106) with multidrug-resistance (3 or more antibiotics from different families), 81% (n=86/106) resistant to ampicillin and 39% (n=41/106) resistant to vancomycin.

### Study of chlorhexidine susceptibility

The minimum inhibitory concentration (MIC) and the minimum bactericidal concentration (MBC) of chlorhexidine gluconate were determined by microdilution (concentration range from 2 to 128 mg/L) using the methodological approach proposed by Clinical and Laboratory Standards Institute methodology for antibiotic susceptibility testing (Mueller-Hinton broth; pH=7.4; 37°C/20h) (54) for the 106 *E. faecium* isolates. The MIC was considered as the first dilution without visible growth after 20h of incubation. The MBC corresponded to the lower chlorhexidine concentration in which 3 or less colonies were detected after the inoculation in Brain Heart Infusion agar with 10 μl (37°C/48h) of each well without visible microbial growth, as described by CLSI for a starting inoculum of 1 x 10^5^ CFU/mL in exponential phase culture (55). *Enterococcus faecalis* ATCC 29212 was used as control strain of chlorhexidine susceptibility, which MIC and MBC were of 16 and 128 mg/L (56, this study), respectively. The assays were repeated two to three times. The assessment of MIC wild type distribution was performed using the ECOFFfinder tool (available at http://www.eucast.org/mic_distributions_and_ecoffs/; ECOFFinder_XL_2010_V2.0) (57), which attempts to fit a log-normal distribution to the presumptive wild type counts by the so-called iterative statistical method. In order to detect differences in chlorhexidine phenotypes distribution among *E. faecium* subpopulations, the ECOFFinder tool was also applied to isolates from clades A1, A2 and B as separated groups. The cutoff of 95% was chosen to set the limits of the wild type population, as recommended by the guidelines of ECOFFinder tool in order to better detect tolerant isolates. The NORM.DIST Excel’s-16.19 function, the estimated mean, the standard deviation and the cumulative normal distribution function option set to TRUE were also used to calculate the probability of occurrence of isolates at the higher concentrations of *E. faecium* population distribution and, consequently, the potential presence of an acquired tolerance mechanism if such probability was too low (57).

As the skin pH ranges between 4 and 6 (39), we also performed a modified microdilution assay using Mueller-Hinton broth adjusted to pH=5 (chlorhexidine-gluconate concentration range: 2-512 mg/L) to evaluate if a more acidic pH than that used in standard conditions (7.4) could have an effect on *E. faecium* susceptibility to chlorhexidine. This assay was performed in 37 whole-genome sequenced *E. faecium* isolates representative of different clades and sources. The remaining experimental conditions as well as the MIC and MBC interpretation were performed as described previously in this section. Aqueous solutions of chlorhexidine-gluconate are most stable within the pH range of 5-8 (https://www.sigmaaldrich.com/content/dam/sigma-aldrich/docs/Sigma/Product_Information_Sheet/c9394pis.pdf). The Mueller-Hinton broth adjusted to pH=5 did not affect the visible growth of *E. faecium* in all strains tested in control wells without chlorhexidine.

### Whole genome sequencing

Among the 106 *E. faecium* studied we selected 37 from different sources [clinical, n=16; healthy human, n=2; long term care patients, n=1; animal production (aquaculture, piggery, aviary), n=6; food of animal origin (bovine meat and milk, poultry carcass, trout), n=9; aquatic environment (river), n=1; wild birds (rook, crow), n=2] and clades (A1, n=19; A2, n=14; B, n=4) to study by whole genome sequence (WGS). Moreover, such genomes corresponded to isolates exhibiting diverse phenotypes to chlorhexidine (MIC ranging from <2 to 32 mg/L). Genomic DNA was extracted (DNeasy Blood and Tissue Kit, Qiagen, Copenhagen, Denmark), with subsequent library construction (Nextera Kit, Illumina, Little Chesterford, UK) and finally WGS (Nextseq, Illumina, Little Chesterford, UK) according to the manufacturer’s instructions to obtain paired-end reads of 2*150 bp in length. Raw sequencing data were analysed using the SerumQC pipeline (https://github.com/ssi-dk/SerumQC) to test the quality of the raw and pre-processed data. The SPAdes genome assembler (58) version 3.10.0 was used for *de novo* assembling the paired-end reads; and SerumQC for evaluating the quality of genome assembly.

The assembled genomes were screened for the presence of genes previously associated with clorhexidine tolerance as *chtR* (coding for conserved DNA-binding response regulator) and *efrE* (codifying for predicted ABC transport system) using BLASTN, and BLASTP analysis was performed to compare the contigs with known sequences of the NCBI database carrying P102H in the ChtR (reference strain was *E. faecium* E1162; GenBank accession no. EFF34003.1; locus_tag EfmE1162_2203) and A290V in the EfrE (reference strain was *E. faecium* 1,231,410; GenBank accession no. EEV54504.1; locus_tag EFTG_02287) proteins, as described for chlorhexidine tolerant *E. faecium* (21, 23). The 37 genomes were also searched for the presence of *qac/smr* genes (*qacA/B, qacEdelta, qacG, qacJ, qacZ*, two *smr*) previously described in *Enterococcus* and other Gram-positive bacteria (27). A database with the *qac* genes sequences was constructed (GenBank accession no. KM083808.1; HG934082.1; NG_048046.1 Y16944.1; HE579074.1; HQ663849.2; GQ900434.1) and genes were tested in our sequenced genomes using the MyDBfinder tool available at Center for Genomic Epidemiology (www.genomicepidemiology.org).

DOOR 2.0 software (http://csbl.bmb.uga.edu/DOOR/) was used to predict the 2CS operon and respective promoter associated with *chtR* and *chtS* genes, previously described by Guzmán-Prieto et al. (21). The information related to each gene product was retrieved from NCBI and PATRIC databases. Comparisons of the amino acid sequences of each operon protein and nucleotide sequences of the predicted promoter were performed with clustalW software (https://www.genome.jp/tools-bin/clustalw) between the reference strain *E. faecium* E1162 and our sequenced *E. faecium* for the identification of specific mutations.

### Comparative genomics

The occurrence of ChtR-P102H and ErfE-A290V mutations was also searched in the *E. faecium* genomes available at the GenBank databases (n=794, last update, 1^st^ December 2018). The ST/clade of these strains was inferred by interrogating the Center of Genomic Epidemiology (www.genomicepidemiology.org) using the in silico genomic MLST tool. Only genomes with a single amino acid change at 102 amino acid on the ChtR protein (H102 or P102) were selected for this analysis. As the EfrE-A290V mutation was only identified in lab mutants (23), we selected all genomes presenting the complete EfrE protein (n=980), which identity was ≥95% with the *E. faecium* 1,231,410 reference strain.

## Data availability

Genome sequences used in this study have been deposited at NCBI under the BioProject accession number PRJNA546230 (https://www.ncbi.nlm.nih.gov/bioproject/?term=PRJNA546230).

## Acknowledgments

The experimental work was supported by the Applied Molecular Biosciences Unit-UCIBIO which is financed by national funds from FCT/MCTES (UID/MULTI/04378/2019). ARF was supported by a post-doc fellowship from REQUIMTE/FCT (DL/57-SFRH/BPD/96148/2013) through Programa Operacional Capital Humano (POCH). All authors from this manuscript do not have any conflict of interest. The funders had no role in study design, data collection and interpretation, or the decision to submit the work for publication.

